# Diclofenac stress responses and biotransformation pathways in the marine diatom *Phaeodactylum tricornutum*

**DOI:** 10.1101/2025.03.13.643036

**Authors:** Leopold Alezra, Emilie Le Floch, Christine Felix, David Rosain, Elena Gomez, Frederique Courant, Eric Fouilland, Giulia Cheloni

**Affiliations:** MARBEC, Univ Montpellier, CNRS, Sète, Ifremer, IRD,France; Master student BEE/EPET- Bidoversité Ecologie Evolution, Sorbonne University; Hydrosciences Montpellier, University of Montpellier, CNRS, IRD, Montpellier, France; Montpellier Alliance for Metabolomics and Metabolism Analysis, Platform on non-target exposomics and metabolomics (PONTEM), Biocampus, CNRS, INSERM, Universit’e de Montpellier, Montpellier, France

## Abstract

Effects of organic contaminants (OCs) on phytoplankton physiology were extensively studied in the last years while a knowledge gap exists regarding the ability of phytoplankton to transform OCs. Knowledge about biotransformation pathways in these organisms lag far behind that of other microorganisms. A better understanding of biotransformation pathways would help identify biomarkers of contaminants exposure, improve microalgae-based water remediation strategies and help better assess contaminants persistence and trophic transfer in natural aquatic environments. The present study investigated diclofenac (DCF) physiological effects, transcriptional responses and metabolism in the marine diatom *Phaeodactylum tricornutum* with the aim of getting an insight on detoxifications pathways. *P. tricornutum* did not result in significant removal capacity of DCF from the exposure medium. Bioconcentration factors varied depending on the exposure concentration (3.9 and 2.7 for 1.5 mg L^-1^ and 10 mg L^-1^ DCF respectively) but remained relatively low. DCF resulted in mild physiological effects on *P. tricornutum* but gene expression analysis indicated that multiple molecular functions and biological processes were altered by DCF exposure. Transcriptomic analysis suggested increased nutrients and energy requirements possibly associated with the contaminant stress and detoxification metabolism. CYP gene expression was not significantly regulated upon DCF exposure but 4’-Hydroxy Diclofenac (OH-DCF), a metabolite generally associated with CYP enzymatic activity, was detected. However, CYP gene expression was not significantly regulated upon DCF exposure. Five additional DCF metabolites with high molecular weight were detected. These metabolites were not previously described in the literature and were suggested to be generated via amino acid (or peptides) conjugation. Gene ontology analysis indicated that amino acid and peptide biosynthetic pathways were regulated upon DCF exposure supporting a possible correlation between organic contaminants detoxification responses and amino acid and protein content in phytoplankton cells. Our findings contribute to highlighting the diversity of biotransformation pathways in phytoplankton and provide mechanistic information about contaminants detoxification.

## 1. Introduction

Millions of tons of synthetic organic chemicals are used annually for industrial, agricultural and consumer’s purposes. These compounds partially find their way to the aquatic environment affecting water quality with consequences for aquatic life (Schwarzenbach et al., 2006). Phytoplankton are aquatic photosynthetic microorganisms that play a key role in the water food chain and global nutrient cycling, they contribute substantially to primary productivity in oceans and are a major sink of carbon dioxide (Doney, 2010). Chemical pollution due to anthropogenic activities may considerably affect phytoplankton. However, phytoplankton exposed to pollutants are not completely unarmed. Adverse outcomes of toxicants are directly linked to the ability of the organism to deal with the toxic insult. When exposed to contaminants, organisms might activate cellular pathways that alter the contaminant intracellular dynamics (toxicokinetic processes) and/or that deal with the stress generated by the chemical exposure (toxicodynamic processes) (EFSA Panel on Plant Protection Products and their Residues (PPR) et al., 2018). Toxicodynamic processes were extensively investigated in phytoplankton but toxicokinetic processes remain underexplored.

Among the different responses that an organism may activate to cope with organic contaminants (OC) exposure, biotransformation plays a determinant role for pollutant fate and transfer (Hellal et al., 2023). Contaminants might be transformed via activation of specific enzymes and the generated transformation products may be either stored in the intracellular environment or released to the extracellular environment. Multiple studies highlighted the promising role that some phytoplankton species have in pharmaceutical removal for biotechnological water treatment processes (Xiong et al., 2018). However, studies focusing on biotransformation pathways in phytoplankton remain scarce (Li et al., 2023). Contaminants biotransformation is investigated in higher aquatic organisms since many years, but in phytoplankton it started to receive attention only recently (Stravs et al., 2017; Ding et al., 2018; Chu et al., 2022; Liakh et al., 2023). What is more, the available studies focused their attention mainly on freshwater cyanobacterial and green algal strains. Information regarding other classes of phytoplankton (e.g marine diatoms) remain underexplored.

In recent years, the release of multiple phytoplankton sequenced genomes allowed the identification of genes encoding for enzymes that may be involved in contaminants biotransformation. Specifically, phytoplankton resulted in multiple copies of cytochrome P450 genes (Teng et al., 2019; Hansen et al., 2021). These enzymes are known to have a prominent role in stress responses and xenobiotic metabolism (Anzenbacher and Anzenbacherová, 2001). CYPs can mediate phase I biotransformation by operating the initial attack of the organic substrate. Members of the CYP family are characterized by a high enzymatic versatility and were reported to be involved in biotransformation of a broad range of different chemicals (Oostenbrink et al., 2012). The role of CYPs in phytoplankton contaminants biotransformation pathways is receiving increasing attention (Li et al., 2023) especially for their potential application in pharmaceutical degradation during biotechnological processes (Xiong et al., 2018; Li et al., 2023). However, multiple aspects related to CYPs contaminant biotransformation remain unknown. CYPs belong to a large family of enzymes that can have very diverse functions. They are involved in synthesis and degradation of multiple compounds related to phytoplankton endogenous metabolism and stress responses (e.g biosynthesis of antioxidant pigments and sterols) rising questions about their specificity in contaminant biotransformation (Li et al., 2023). In phytoplankton, there is no agreement on which sub-family of CYPs enzymes is involved in contaminants metabolism. What is more, these enzymes are known to form enzymatic complexes with redox partners for their metabolic function. At present, partners required to form the active complex for efficient contaminant biotransformation in phytoplankton are unknown (Li et al., 2023).

The present paper investigates the physiological effects, transcriptional responses and biotransformation pathways of diclofenac (DCF) in the marine diatom *Phaeodactylum tricornutum*. A special attention was given to the contribution of CYPs that were proposed to be involved in the generation of phase I metabolites in multiple organisms (Prior et al., 2010; Fu et al., 2017; Ben Ouada et al., 2019; Fu et al., 2020; Liakh et al., 2023). DCF is reported to be efficiently removed from exposure medium by phytoplankton (Escapa et al., 2016; Ben Ouada et al., 2019; Hifney et al., 2021) and to induce physiological effects at concentrations above the mg L^-1^ range (Bácsi et al., 2016; Majewska et al., 2018; Ben Ouada et al., 2019; Harshkova et al., 2021; Sharma et al., 2023). We hypothesized that effects observed at such high DCF concentration may be associated with the ability in phytoplankton to detoxify this compound and that CYPs play a determinant role in this process.

## 2. Materials and Methods

### 2.1 Algae culturing and Diclofenac exposure experiments

Cultures of *Phaeodactylum tricornutum* (CCAP1055/18, axenic strain, SAMS Culture Collection of Algae and Protozoa) were grown in glass Erlenmeyer flasks with f/2+Si medium (Guillard and Ryther, 1962) under controlled conditions (20°C, continuous agitation at 120 rpm, warm white fluorescent lighting at 50 μE with a 16/8 light:dark cycle). A growth curve of the CCAP1055/18 strain under the described culturing conditions can be found in supplementary information (Figure S1). Axenicity of *P. tricornutum* cultures was assessed in the beginning and at the end of the experiments. Samples of cultures were collected and stained with the SYTO BC bacteria stain (Bacteria counting kit, Molecular Probes, Invitrogen) following the manufacturer protocol. Samples were analyzed using the flow cytometer BD Accuri C6 plus (BD biosciences). Details on gating strategy and results are available as supplementary material (Figure S2).

### 2.2 Chemicals and analytical standards for MS analysis

The analytical standard of Diclofenac (DCF), Diclofenac-d4 (DCF-d4), 4’-Hydroxy Diclofenac (OH-DCF) and 4’-Hydroxy Diclofenac-d4 were purchased from LGC Standards. All standards were purchased at analytical grade (purity >98%). Stock standard solutions of each compound were prepared in methanol at a concentration of 1 mg/mL. All the standard solutions were stored at 20°C. A working standard solution for each standard was prepared at concentration of 1 μg/mL in methanol. Ultrapure water was generated by a Millipore Milli-Q system (Milford, MA, USA). Methanol (MeOH), and HPLC-grade acetonitrile (ACN) were obtained by Carlo Erba (Val de Reuil, France). Formic acid (FA) (purity, 98%) was supplied from Fisher Scientific Labosi (Elancourt, France). Solid phase extraction (SPE) Oasis HLB™ cartridge (60mg, 3cc) was purchased from Waters, (Milford, MA, USA).

### 2.3 Diclofenac exposure experiments

*P. tricornutum* cultures in the exponential phase were collected and diluted in order to reach an initial cell density of 1×10^6^ cell/mL. Cells were inoculated in f/2+Si medium (control cultures) or in f/2+Si medium spiked with the amount of diclofenac stock solution required to reach the desired final concentration of the contaminant. Diclofenac sodium salt (purity ≥ 98%, Sigma-Aldrich, Steinheim, Germany) stock solutions were prepared right before the exposure experiments by dissolving product powder in f/2+Si medium without addition of solvent. Solutions were sonicated three times for 5 minutes in an ultrasonic bath (37 kHz frequency), let equilibrate for 30 minutes and filtered on a 0.22 μm sterile filtering unit before utilization. DCF concentrations in the exposure medium were determined at the beginning and at the end of each exposure test. Stability of DCF in exposure medium was verified by preparing control Erlenmeyer flasks containing f/2 medium spiked with DCF but not inoculated with algae, incubated under the same conditions as cultures. DCF concentrations resulted to be stable in the exposure medium along the 72 hours duration of the test (Figure S3B).

Control and exposed cultures were incubated for 72 hours under the controlled conditions previously described. Experiment starting time was set in the middle of the light phase.

Three different DCF exposure experiments were performed. The first experiment aimed at testing effects of different DCF concentrations (ranging between 0.5 and 50 mg L^-1^) on algal growth.

The second experiment aimed to investigate physiological effects and gene expression. It included two selected DCF concentrations (1.5 and 10 mg L^-1^) and unexposed cultures. The concentration of 1.5 mg L^-1^ and 10 mg L^-1^ were selected for further investigation as they represented the treatments at which the maximal and minimal growth values were observed (Figure 1). Experiments were run in triplicates starting from independent cultures.

**Figure 1:**
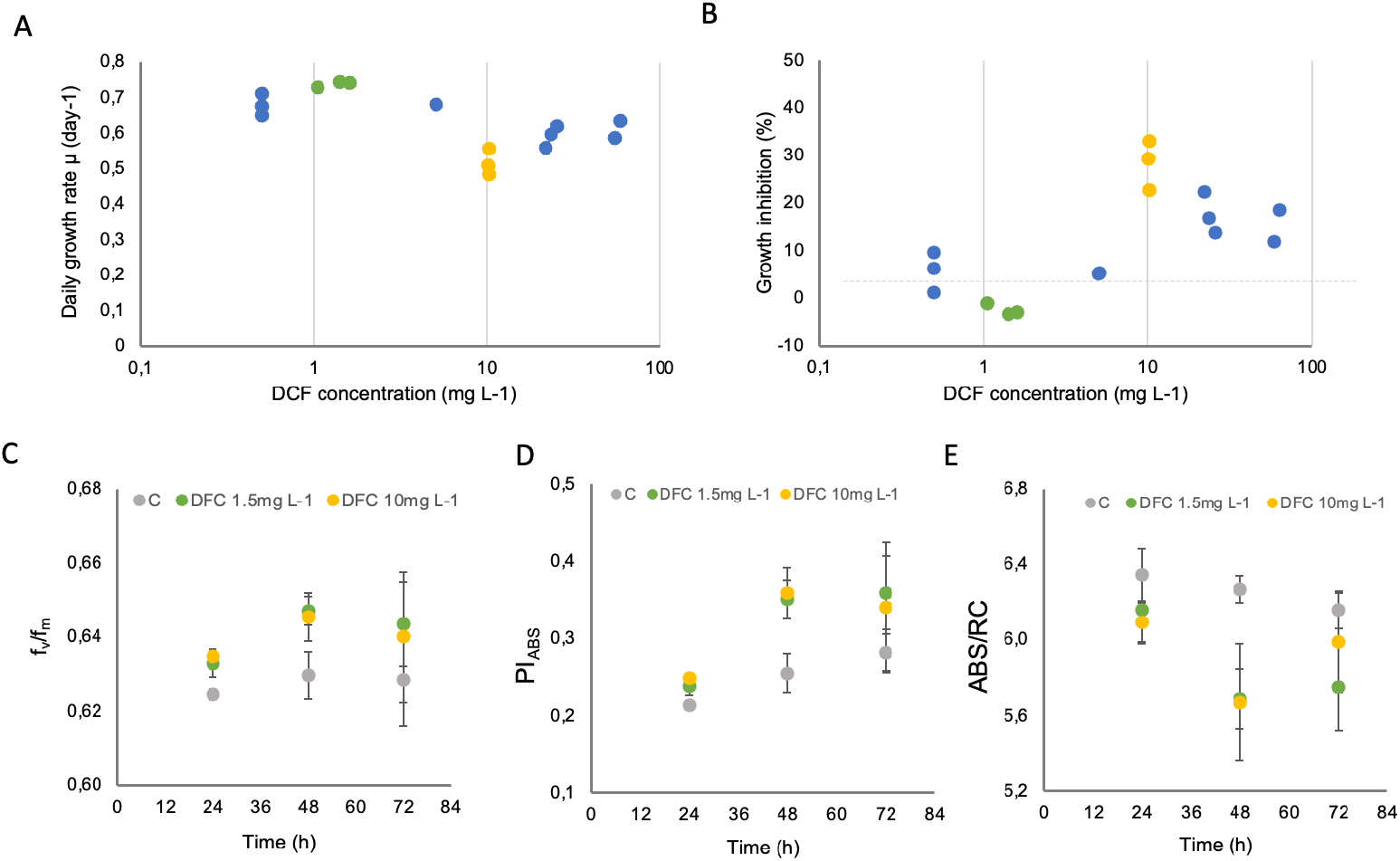
Effects of DCF on *P. tricornutum* growth and photosynthetic activity. (**A**) Daily growth rate measured in *P. tricornutum* cultures upon 72 hours exposure to different DCF concentration expressed as measured concentrations. Daily growth rate in control cultures was 0,72±0,01. (**B**) Growth inhibition observed upon 72 hours exposure to DCF expressed as percentage with respect to unexposed control cultures. Green and yellow dots correspond to nominal concentrations (1.5 mg L^-1^ and 10 mg L^-1^ respectively) that were selected for subsequent tests, values of measured concentrations are available in Figure S3B. (**C**) Fv/Fm values measured for *P. tricornutum* cultures after 72 hours exposure to 1.5 mg L^-1^ DCF (green dots) and 10 mg L^-1^ DCF (yellow dots) or unexposed control cultures (grey dots). (**D**) PI_ABS_ values measured for *P. tricornutum* cultures after 72 hours exposure to 1.5 mg L^-1^ DCF (green dots) and 10 mg L^-1^ DCF (yellow dots) or unexposed control cultures (grey dots). (**D**) ABS/RC values measured for *P. tricornutum* cultures after 72 hours exposure to 1.5 mg L^-1^ DCF (green dots) and 10 mg L^-1^ DCF (yellow dots) or unexposed control cultures (grey dots). In plots C, D and E results are presented as average values of three independent replicates.

The third experiment aimed at producing samples for LC-MS investigation and included unexposed cultures, two DCF concentrations (1.5 and 10 mg L^-1^) and a treatment combining 10 mg L^-1^ DCF and CYP inhibitor Fluconazole. Fluconazole is a CYP51 inhibitor that was reported to be effective between 10 and 200 mg L^-1^ without having effects on *P. tricornutum* growth (Fabris et al., 2014). Fluconazole (purity ≥ 98%, Sigma-Aldrich, Steinheim, Germany) stock solutions were prepared in f/2+Si medium without addition of any solvent following the procedure described for DCF stock solutions. Fluconazole concentrations in the exposure experiment corresponded to 21.9±2.2 mg L^-1^ and were directly quantified via HPLC-UV analysis using the same method adopted for DCF. These concentrations did not have significant effects on *P. tricornutum* growth, photosynthetic efficiency and intracellular oxidative stress. Experiments were run in five replicates starting from independent cultures. For all experimental treatments, cells were harvested After 72h exposure by centrifugation at 5000 g for 10 minutes. Supernatant was discharged and pellets were rinsed with 1 mL fresh f/2+Si medium, quickly resuspended, transferred in a 2 mL tube and centrifuged again at 10000 g for 5 minutes. Supernatant was discharged, pellets were flash frozen in liquid nitrogen and stored at -80°C before lyophilization.

### 2.4 Diclofenac measurements in exposure medium

Aliquots of 5 mL of test medium (inoculated or not with *P. tricornutum*) were collected, filtered on a 0.22 μm sterile filtering unit and stored at -20°C until analysis. DCF concentrations in the exposure medium were quantified via HPLC-UV. Aliquots of 2 mL of each sample were loaded on solid-phase extraction (SPE) cartridges (Oasis HLB 60mg 3cc) that were preconditioned with 3mL MeOH and 3 mL MilliQ water. Columns were rinsed with 4 mL deionized water and dried. Components were eluted with 4 mL MeOH, dried under nitrogen stream and resuspended in 200 μL (acetonitrile:water) used for HPLC-UV analyses. HPLC analysis was performed with a Vanquish HPLC (ThermoFisher Scientific, Bremen, Germany) equipped with an Vanquish UV detector (ThermoFisher Scientific, Bremen, Germany) and a Luna 5μm C18 (150×4,6 mm) capillary column (Phenomenex, Torrance, CA, USA). Detailed information about the HPLC method used to analyze the samples can be found in the supporting information (Figure S3A).

### 2.5 Characterization of physiological effects

Aliquots of cultures were collected daily (time 0, 24, 48 and 72 hours) in order to investigate multiple physiological parameters. Growth was measured via direct cell counting after Lugol fixation, 5% rose bengal staining and appropriate dilution in the culture medium, using a Luna II automated cell counter (Logos Biosystems, division of Aligned Genetics, Anyang, South Corea). Changes in photosystem II activity (Fv/Fm, ABS/RC and PI_ABS_) were determined using an AquaPen-C AP-C 100 fluorometer (Photon System Instruments, Drásov, Czech Republic) using OJIP chlorophyll fluorescence induction kinetics protocol (Strasser et al, 2000, Stirbet and Govindjee, 2011). Cultures were collected and incubated 30 minutes in the dark before analysis. When required, cultures were diluted in their own exposure medium obtained by filtering an aliquot of culture on a 0.22 μm sterile cellulose acetate filtering unit. Intracellular oxidative stress was measured via flow cytometry upon staining with CellROX™ green flow cytometry assay kit (Molecular Probes, Life Technologies). Information about the adopted procedure can be found in the supplementary information document (Figure S4).

### 2.6 Transcriptomic analysis

Samples for transcriptomic analysis were collected at the end of the DCF exposure experiments (72 hours incubation). Cultures were filtered onto 2 polycarbonate filters (0.22 um) in order to harvest at least 2×10^8^ cells using a vacuum filtering system. Filters were washed with 10 mL of f/2+Si medium and collected in cryotubes that were flash frozen in liquid nitrogen and stored at -80°C. Total RNA was extracted using TriPure isolation reagent (ref 11667157, Roche,) following the manufacturer protocol. Extracted RNA was treated with DNase RQ1, RNAse free (Promega) and further cleaned up using RNeasy Mini kit columns (Qiagen, Gemrmany). Integrity and concentration of extracted RNA was verified using Nanodrop (and Qubit RNA broad range kit (ThermoFisher Scientific, Bremen, Germany). RNA sample quality control was performed using a TapeStation system (Agilent). RNA libraries were constructed using the Illumina TruSeq stranded mRNA Sample Preparation kit (Illumina, USA) and sequenced with Illumina Hiseq 2000 platform. Analyses were successfully performed on all of 125 bp/150 bp paired-ends samples. Trimmed reads were mapped to the reference genome with HISAT2 (Kim et al., 2015). After the read mapping, StringTie (Pertea et al., 2015) was used for transcript assembly. Expression profile was calculated for each sample and transcript/gene as read count, FPKM (Fragment per Kilobase of transcript per Million mapped reads) and TPM (Transcripts Per Kilobase Million). DEG (Differentially Expressed Genes) analysis was performed on 3 comparisons pairs (1.5 mg L^-1^ DCF treatment versus Control; 10 mg L^-1^ DCF treatment versus Control; 10 mg L^-1^ DCF treatment versus 1.5 mg L^-1^ DCF treatment) using DESeq2 (Love et al., 2014). The g:Profiler (Kolberg et al., 2023) tool was used to perform gene ontology (GO) enrichment analysis on DEG genes, a significance threshold of 0.05 was applied.

### 2.7 Diclofenac and biotransformation products analysis in phytoplankton

#### 2.7.1 Sample preparation of extracts

15 mg of the freeze-dried biomass samples were ground using a Bullet Blender Gold (Next Advance, USA) at power 10 for 5 minutes, spiked with 10 ng/mg of DCF-d4 and 0,1 ng/mg of OH-DCF-d4 standards and extracted twice using 1 mL of acetonitrile (ACN) containing 0.4% formic acid. In order to ensure homogeneous extractions samples were blended for additional 2 minutes at power 10 and sonicated in an ultrasound bath for 5 minutes at 37 kHz frequency. Supernatants were collected after 5 minutes centrifugation at 10000 g. Extracted compounds were evaporated to dryness and resuspended in 100 μL MeOH. Extracts were diluted with 3 mL MilliQ water and purified using solid-phase extraction (SPE) cartridges (Oasis HLB 60mg, 3cc) that were preconditioned with 2 mL MeOH and 2 mL MilliQ water. Cartridges were rinsed with 2mL MeOH/H_2_O (5/95) and dried. Components were eluted with 3 mL MeOH, dried under nitrogen stream and resuspended in 200 μL ACN/H_2_0 (20/80) before filtering on 0.2 μm PTFE filters (Fischer Scientific, Bremen, Germany).

To perform DCF quantification, an extracted calibration curve was prepared in the same way of extracts with samples spiked at 1, 5, 10, 25 and 50 ng/mg of DCF (DCF-d4 at 10 ng/mg in each calibration sample) and 0.01, 0.05, 0.1, 0.25, 0.5 and 1 ng/mg of OH-DCF (4-OH-DCF-d4 at 0.01 ng/mg in each calibration sample). R^2^ values were above 0.99 for all target analytes. To determine DCF bioconcentration in phytoplankton, a bioconcentration factor (BCF) was calculated at 72h as follows:

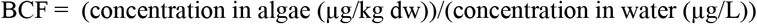

#### 2.7.2 HPLC-HRMS Analysis

The injections were performed on a Q Exactive Orbitrap LC-HRMS (ThermoFisher Scientific, Bremen, Germany) equipped with a heated electrospray ionization source (HESI). A reverse phase pentafluorophenylpropyl (PFPP) analytical column (100 × 2.1 mm; 3 μm particle size; Sigma Aldrich) was used for LC separation. The LC mobile phases were water (A) and ACN (B), both modified with 0.1% formic acid. Each sample (10 μL) was loaded onto the column. The flow rate was 250 μL/min according to the following gradient (A/B): 95/5 at 0 min, 86/14 at 3 min, 66/34 at 14 min, 55/45 at 18 min, 5/95 from 20 to 25 min and a return to the initial conditions at 28 min, followed by a 7 min re-equilibration period (95/5), for a total run time of 35 min. Samples were analyzed in negative electrospray ionization modes (ESI-). ESI-parameters were as follows: sheath gas 55 arbitrary units (AU), auxiliary gas 10 AU, capillary temperature 300 °C, heater temperature 250°C, while the electrospray voltage was set at -3 kV. The S-lens RF level was set at 95. Full scan mode with a mass range of *m/z* 100–750 at a mass resolving power of 35,000 was used.

#### 2.7.3 Biotransformation products identification strategy

Chemical profiles from control, DCF 1 and 10 mg exposed samples, and DCF+Fluconazole exposed samples (5/condition) were generated by HPLC-HRMS (§2.5.2). The presence of DCF metabolites was assessed by a non-targeted approach. Raw data were converted into mzXML files with MSConvert freeware (ProteoWizard 3.0, Holman et al., 2014). ESI-acquisitions were processed using the XCMS package (Smith et al., 2006) in the R environment to integrate the chromatographic peaks per sample. This multi-step strategy has already been described by Bonnefille et al. (2017). XCMS parameters were applied as follows: the *m/z* interval was set at 0.01, the signal to noise ratio threshold at 10, the group bandwidth at 8, and the minimum fraction at 0.5. After data processing, XCMS generated a table containing peak information and feature abundances per sample. The peaks detected only in the exposed samples (absent in the control samples) and found in at least three replicates among the five were considered to indicate potential DCF metabolites. Extracted ion chromatograms (EICs) were also extracted to confirm the absence of signal in the controls. Elemental compositions of unknown metabolites were generated by the Thermo Xcalibur Qual Browser (XCalibur 4.2.47), and those with the most suitable C, H, N, O and Cl compositions, based on the DCF composition, were reported. Metabolites presented a specific chlorinated/dichlorinated isotopic pattern of M/M+2/M+4 reflecting the natural abundance of Cl isotopes (^35^Cl 76% and ^37^Cl 24%).

### 2.8 Statistical analysis

Statistical differences were tested comparing treatments versus control or treatments with different DCF concnetrations using two-sample test Mann-Whitney. Significant differences were identified using a criterion of a p < 0.05

## 3. Results

### 3.1 Physiological effects upon DCF exposure

#### 3.1.1 Growth inhibition upon DCF exposure

DCF had no significant impact on growth for concentrations ranging between 0.5 and 5 mg L^- 1^ (Figure 1A). For the 1.5 mg L^-1^ treatment a slight (not significant) increase in growth was observed suggesting a possible hormetic effect of this chemical compound. The maximal growth inhibition (25% on average) was observed under 10 mg L^-1^ DFC exposure (Figure 1B) followed by a slight decrease in the effect at higher concentrations.

#### 3.1.2 Effects on photosynthetic activity and oxidative stress

At concentrations of 1.5 and 10 mg L^-1^, DCF had no significant effects on photosynthetic activity (fv/fm) (figure 1C). An effect (although not statistically significant) was observed on both concentrations after 48h for PI_ABS_ parameter (performance index for energy conservation from photons absorbed per PSII to the reduction of Qb) and ABS/RC (the average absorbed photon flux per PSII reaction center, apparent antenna size of an active PSII). No significant effect of DCF was observed on intracellular oxidative stress (Figure S4).

### 3.2 Gene expression upon DCF exposure

The number of genes differentially expressed upon 1.5 mg L^-1^ and 10 mg L^-1^ treatments with respect to control cultures were 1331 and 1598 respectively (Figure 2A). In both treatments the number of up-regulated genes was higher (933 and 925 genes for 1.5 mg L^-1^ and 10 mg L^-1^ treatments respectively) than the number of down-regulated genes (398 and 673 genes for 1.5 mg L^-1^ and 10 mg L^-1^ treatments respectively). The 1.5 mg L^-1^ and 10 mg L^-1^ treatments shared 553 genes differentially expressed with respect to control (Figure 2B) but more than the half of the differentially expressed genes in each treatment were specific to the DCF exposure concentration. Functional analysis of genes differentially expressed in the 10 mg L^-1^ treatment with respect to 1.5 mg L^-1^ treatment detected only one significant molecular function associated to DNA-binding transcription factors (Figure 2C). These genes were significantly down regulated in the 10 mg L^-1^ treatment (Table S1).

**Figure 2:**
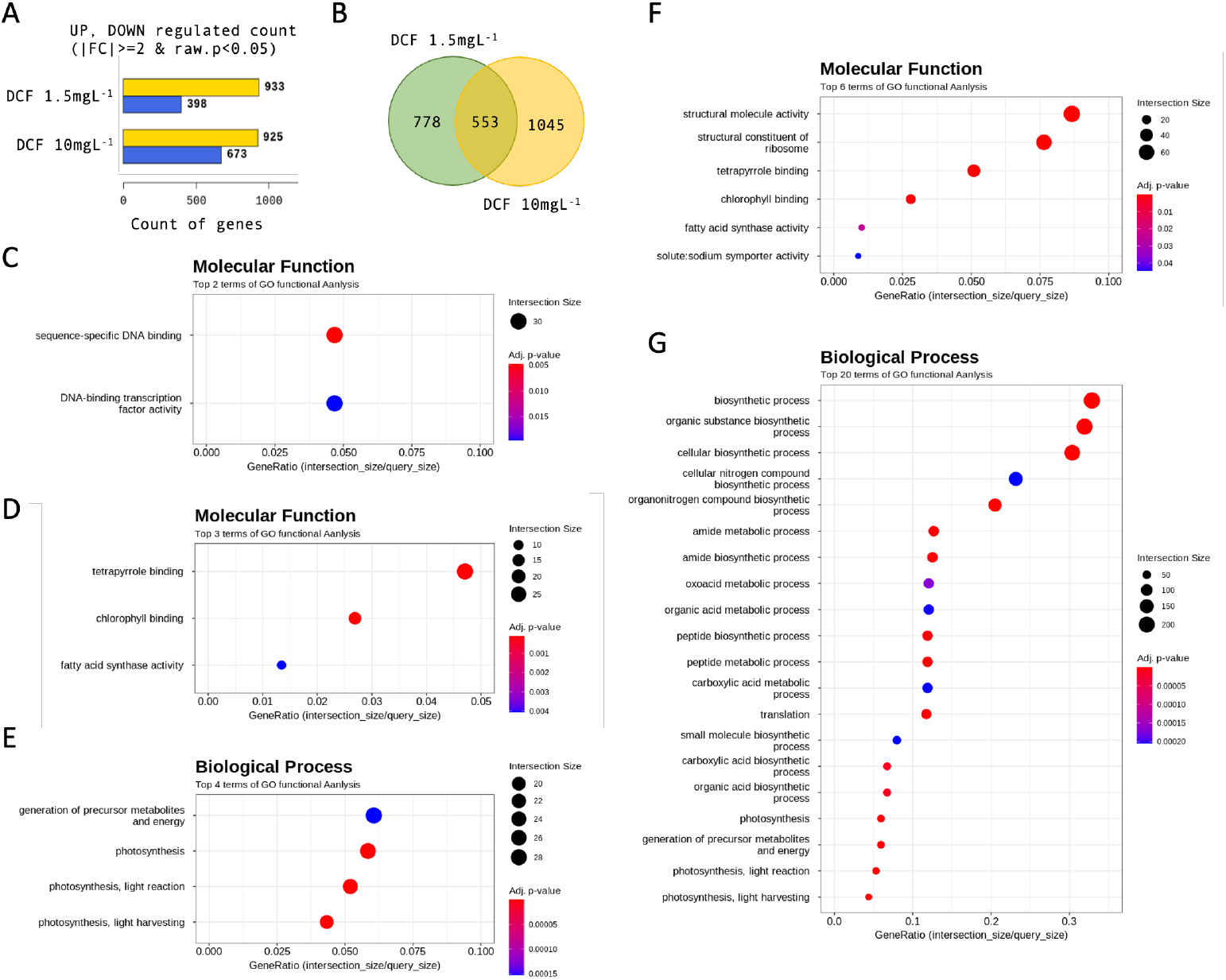
Comparative analysis of gene expression in *P. tricornutum* cultures treated for 72 hours with 1.5 mg L^-1^ and 10 mg L^-1^ DCF concentrations with respect to untreated control. (A) Number of up regulated (yellow bar) and down regulated (blue bar) genes in DCF treated cultures. (B) Venn diagram of differentially expressed genes observed in 1.5 mg L^-1^ (green set) and 10 mg L^-1^ (yellow set) DCF treatments. (C) Scattered plot of GO function (molecular function) enrichment in and 10 mg L^-1^ DCF treatment with respect to 1.5 mg L^-1^ treatment. (D) Scattered plot of GO function (molecular function) enrichment in and 1.5 mg L^-1^ DCF treatment with respect control. (E) Scattered plot of GO function (biological process) enrichment in and 1.5 mg L^-1^ DCF treatment with respect control. (F) Scattered plot of GO function (molecular function) enrichment in and 10 mg L^-1^ DCF treatment with respect control. (G) Scattered plot of GO function (biological process) enrichment in and 10 mg L^-1^ DCF treatment with respect control.

Functional analysis of differentially expressed genes in 10 mg L^-1^ treatment with respect to control indicated that genes involved in structural constituents of ribosome were significantly regulated (Figure 2F) as well as genes involved in multiple biological processes for amide and peptides biosynthetic processes and for translation (Figure 2G). These functions were not significantly regulated in the 1.5 mg L^-1^ treatment. On the other hand, 1.5 mg L^-1^ treatment and 10 mg L^-1^ treatment had in common the regulation of genes with tetrapyrrole binding function (heme binding and chlorophyll binding (Table S2)) and fatty acid synthase activity (Figure2 D and F) and the regulation of genes involved in photosynthetic processes (light reaction and light harvesting) (Figure2 E and G).

Gene ontology enrichment analysis of differentially expressed genes in 1.5 mg L^-1^ and 10 mg L^-1^ treatments did not highlight significant regulation of genes involved in Iron homeostasis. However, ISIP genes (involved in iron internalization and trafficking (Kazamia et al., 2022)) and CREG1 (proposed to be involved in phytotransferrin endocytosis (Turnšek et al., 2021)) resulted to be among the top 15 up regulated genes in DCF treatments (Table S3). These results are coherent with the fact that multiple genes encoding for hemoproteins were significantly upregulated upon DCF exposure (Figure 3).

**Figure 3:**
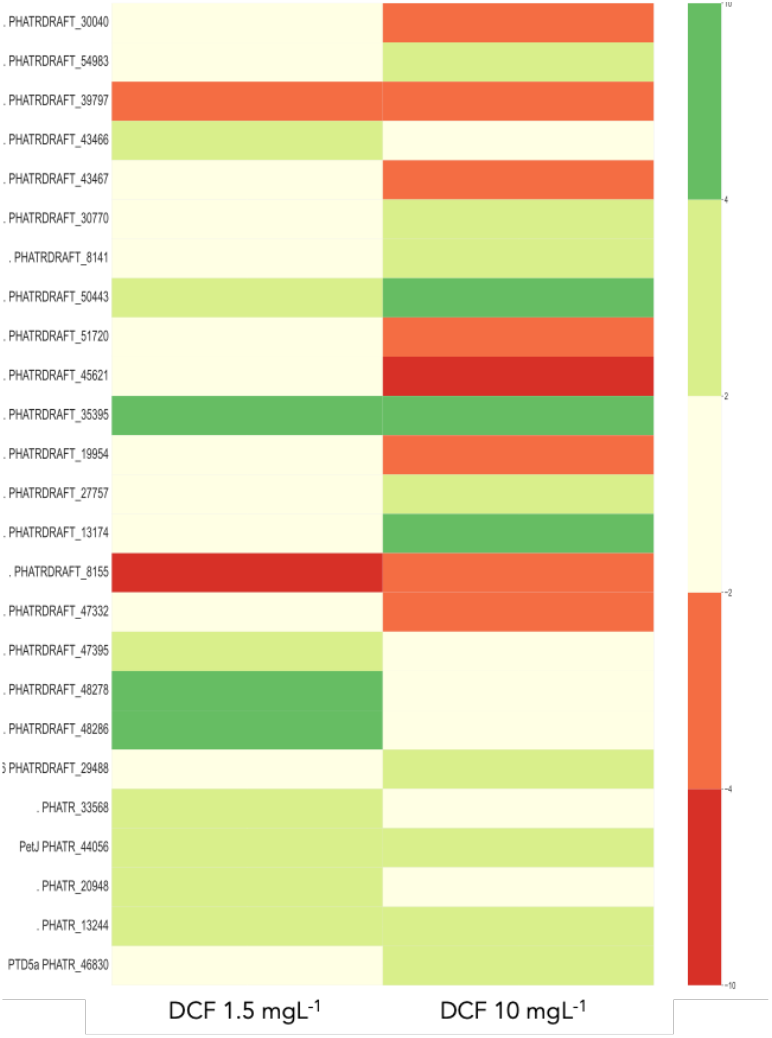
Heat map of fold change values obtained for transcripts of hemeproteins that were significantly upregulated in the 1.5 mg L^-1^ treatment, in the 10 mg L^-1^ treatment or in both treatments. Color code indicates intervals of fold change (dark red values between -4 and -10; light red -4 and -2; white -2 and 2; light green 2 and 4, dark green 4 and 10).

Among the genes encoding for hemeproteins, those with peroxidase activity were generally upregulated (Figure 3 and Table S2). On the other hand, among the CYP genes identified in *P. tricornutum*, only two were differentially expressed and CYP5165A2 (PHATRDRAFT_43466) was upregulated in the 1.5 mg L^-1^ treatment whereas CYP5165A1 (PHATRDRAFT_43467) was downregulated in 10 mg L^-1^ treatment (Table S2). Transcripts of multiple hemeproteins with cytochrome B5-like domain were differentially expressed upon DCF treatment (Figure 3 and Table S2).

### 3.3 Measurements of DCF and OH-DCF

DCF concentrations in the exposure medium did not change significantly through time neither in the 10 mg L^-1^ treatment (Figure 4A) nor in the 1.5 mg L^-1^ treatment (Figure 4B). On the other hand, significant difference in the intracellular DCF contents was measured between the two treatments (Figure 4C). Higher amounts of intracellular DCF were measured in cells that underwent the treatment with the higher DCF concentration. BCF resulted to be 3,93 and 2.68 for the treatment at 1.5 mg L^-1^ and 10 mg L^-1^ treatment respectively.

**Figure 4:**
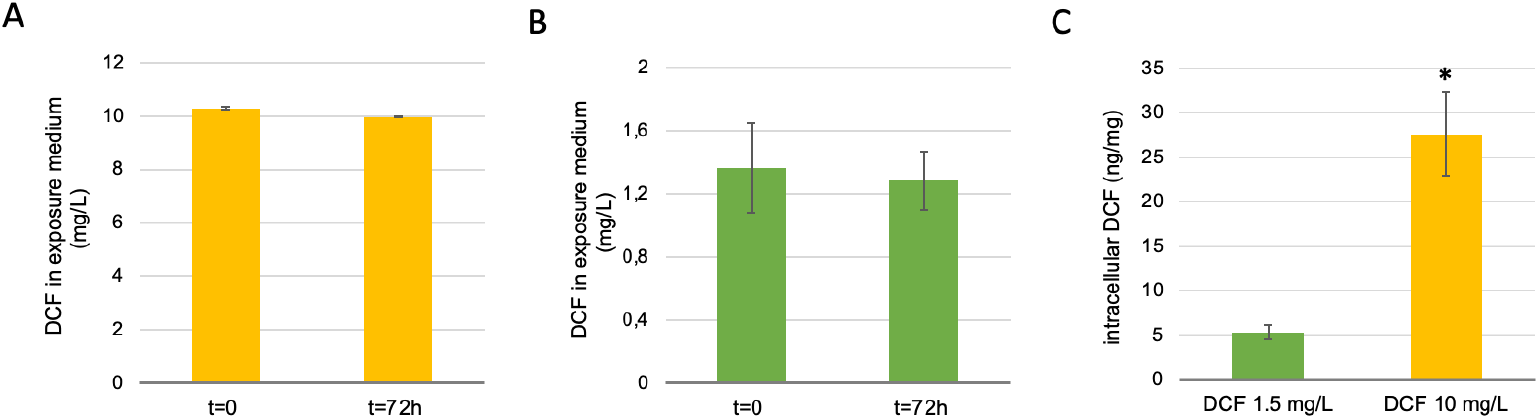
Quantification of DCF in exposure medium at the beginning and in the end (72h) of exposure tests for the 10 mg L^-1^ (A) and 1.5 mg L^-1^ (B) concentrations. (C) Intracellular DCF content is measured for cultures exposed to 1.5 mg L^-1^ (green bar) and 10 mg L^-1^ (yellow bar). Values are presented as average ± st. dev. * indicate a significant difference (p<0,05).

### 3.4 DCF metabolites and transformation pathways

Six different DCF metabolites (M2-M7) were detected in cells treated with 10 mg L^-1^ DCF (Table 1) and absent from the controls. Only two of them (M3 and M4) were observed in cells treated with 1.5 mg L^-1^ DCF suggesting that either the other metabolites are not generated or their amount is lower than the detection limit of the technique.

**Table 1.**
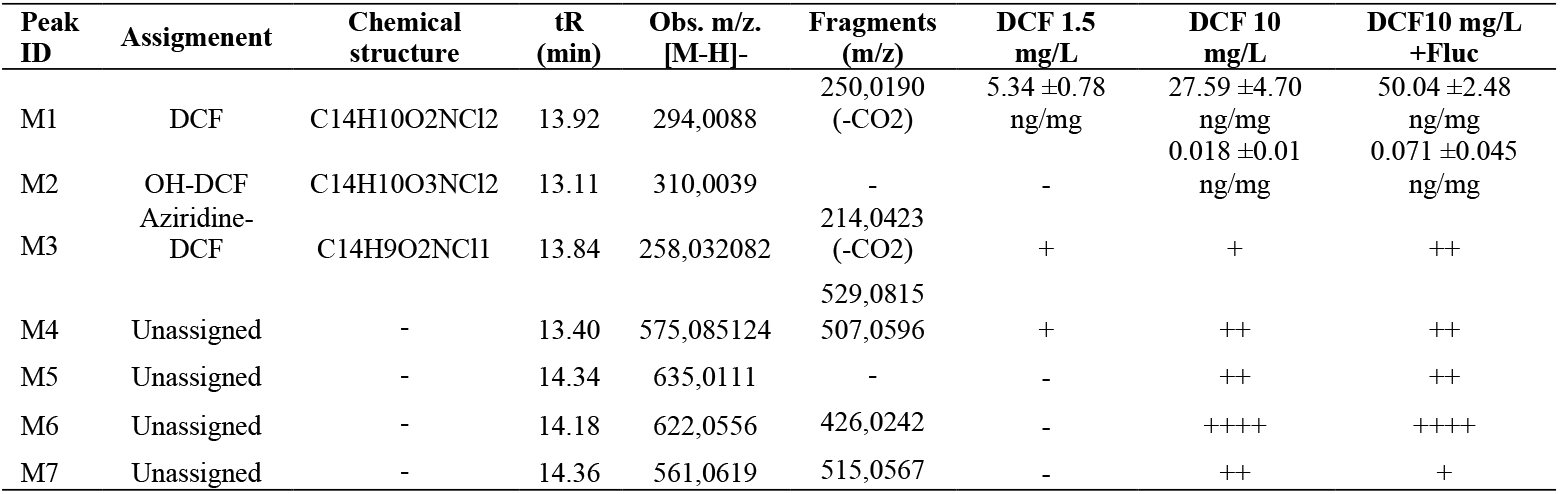
DCF metabolites detected via HPLC-MS analysis (+ signs indicate the abundance of peaks measured for the metabolite)

Among the metabolites observed in the 10 mg L^-1^ treatment we could detect 4’ Hydroxy-diclofenac (OH-DCF, M2, confirmed with the analytical standard) even if amounts of this metabolite were very low. M3 was detected in cells treated with 1.5 and 10 mg L^-1^ DCF and based on its mass we deduce that this metabolite underwent loss of one atom of H and one atom of Cl. For the other metabolites (M4-M7) it was not possible to propose a structure as they did not correspond to DCF metabolites previously reported in the literature. All of them (M4-M7) are characterized by an atypically large mass that may be associated with phase II modifications of different groups. The most abundant metabolite observed in the 10 mg L^-1^ treatment was M6 whose peak intensity was higher than the one observed for DCF (Figure S5).

Upon addition of CYP inhibitor (fluconazole), the amount of intracellular DCF increased significantly while concentrations of OH-DCF remained unchanged (Table 1). Changes were observed as well for M3 and M7 metabolites whose peak intensity increased and decreased respectively (Figure S6).

## 4. Discussion

Dichlorofenac (DCF) inhibited the growth of *P. tricornutum* by 25% at 10 mg L^-1^. At the lowest concentrations tested, DCF had a positive effect on growth (although not significant). Hormetic effect of similar ranges of concentrations were previously reported for *Parachlorella kessleri* (Jiménez‐Bambague et al., 2022) and for *Picocystis* sp. (Ben Ouada et al., 2019). The dose response growth inhibition curve did not follow the classical sigmoidal function previously reported for DCF in *Chlamydomonas reinhardtii* (Majewska et al., 2018). However, in our experiments, DCF treatments were limited to lower concentrations with respect to those tested for *C. reinhardtii*. In our tests the highest DCF concentration was 50 mg L^-1^. This concentration is close to the solubility limit expected for DCF in aqueous solutions with high salt concentrations such as f/2 medium.

The main mode of toxic action described for DCF is the inhibition of prostaglandin synthesis via inhibition of cyclooxygenase-1 (COX-1) and cyclooxygenase-2 (COX-2) (Gan, 2010). Marine diatoms have COX enzymes for prostaglandin synthesis (Di Dato et al., 2017) but COX genes are absent on Phaeodactylum genome (Di Dato et al., 2020). DCF mode of toxic action was investigated in *Chlamydomonas reinhardtii* and was reported to be connected to photosynthesis inhibition associated with the “silencing” of a fraction of the PSII reaction centers (Majewska et al., 2018). In this work we analyzed multiple photosynthetic activity endpoints that were reported to be affected by DCF in *C. reinhardtii*. However, significant effects were not observed after 72 hours incubation. Effects observed after 48 h treatment on Performance Index (PI_abs_ which quantifies the overall functionality of the electron flow through PSII) and ABS/RC (average absorbed photon flux per reaction center) were compensated after 72h treatment. The recovery effect is consistent with transcriptomic data and gene ontology enrichment analysis that indicated a significant upregulation of genes associated with photosynthesis, light reaction and light harvesting. Most of the genes encoding for light harvesting complexes (LHC) were upregulated upon DCF treatment.

DCF was reported to increase antioxidant enzyme activity in green algae (Majewska et al., 2018; Sharma et al., 2023) but no effects on intracellular oxidative stress were observed in our tests. Nevertheless, genes encoding for enzymes involved in oxidative stress responses were up regulated upon DCF treatment suggesting that the antioxidant response may effectively compensate for the possible generation of ROS.

Even if DCF did not substantially affect *P. tricornutum* physiological parameters, a significant change in gene expression was observed upon DCF treatment. More than a thousand genes were differentially expressed out of the 12,089 predicted in the organism genome (Phatr3 genome reannotation (Rastogi et al., 2018)). To our knowledge, this is the only work that explored gene expression upon DCF treatment in phytoplankton and no information is available for comparative discussion. However, transcriptomic information is available for *P. tricornutum* exposed to the contaminant naphthenic acid (Zhang et al., 2021). 96h exposure with naphthenic acid resulted in the enrichment of molecular functions and biological processes that were observed as well upon DCF treatment in the present study. Naphthenic acid affected photosynthesis-related genes (that were up-regulated), tetrapyrrole binding proteins and tetrapyrrole related metabolic processes. Also, enrichment in ribosome biogenesis, ribonucleoprotein complex biogenesis and biosynthesis of amino acid process was observed. Such similarity in the transcriptomic responses suggest that DCF and naphthenic acid have a similar mode of toxic action.

In our work, genes that resulted to be differentially expressed upon DCF treatments are related to nutrient starvation. Nitrate reductase (Phatr3_J54983) was up regulated as well as genes involved in iron homeostasis (ISIP2B, ISIP2A, ISIP3, CREG1, FRE2, FRE3 and ZIP1). Cultures at the cell density measured after 72 hours incubation are not expected to be starved in f/2 medium as shown in Fig S1. The observed effects may be associated with higher nutrients and energy requirements to cope with contaminants and detoxification metabolism. For instance higher iron quotas may be necessary to sustain the synthesis of hemoproteins whose genes were shown to be up-regulated. Heme-containing proteins (notably CYPs) are involved in a large variety of key functions in photosynthetic organisms, including stress responses.

No significant changes in CYP gene expression was observed upon DCF treatment despite the fact that CYP associated DCF metabolites were detected (OH-DCF). CYP genes may not be differentially expressed upon contaminant stress in diatoms or CYP activity may be modulated through their redox partners. *P. tricornutum* has one gene for Nadph-cytochrome P450 reductase (CPR, Phatr3_J45758) and multiple genes encoding for proteins with Cytochromes B5 like domains. CPR gene was not differentially expressed upon DCF treatment. However, this gene may undergo alternative regulation mechanisms. For instance, CPR gene expression was associated with microRNA regulation in *C. reinhardtii* (Gao et al., 2016). Two genes with Cytochromes B5 like domains (PHATRDRAFT_35395 and PHATRDRAFT_50443) were over expressed upon DCF treatment and their levels of expression increased with DCF exposure concentration. PHATRDRAFT_50443 (also known as EG02619) is considered to be a putative D6 fatty acid desaturase but this role was not confirmed by functional genomic studies (Dolch et al., 2017). CYP activity may be regulated via expression of specific Cyt B5 redox partners.

The amount of DCF accumulated in *P. tricornutum* cells after 72h incubation was relatively low with respect to exposure concentrations suggesting that either this chemical compound is taken up at low rates from the extracellular environment or it is efficiently transformed. The phase I metabolite OH-DCF was detected in *P. tricornutum* cells exposed to DCF but its concentration was considerably low (1000 fold lower than the parent compound) suggesting that OH-DCF is used as substrate for Phase II modifications. Multiple metabolites with high molecular weight were identified after 72h incubations, some of which in an amount that was estimated to be higher than the one measured for DCF. The low intracellular content of Phase I metabolites coupled with the higher abundance of Phase II metabolites indicate that biotransformation pathways are shifted toward end products. In phytoplankton, abundance of Phase I and Phase II metabolites was reported to change through incubation time with multiple end products accumulated in the long term in the intracellular environments while intermediate products accumulated in the exposure medium (Liakh et al., 2023). Metabolites observed in this study may represent only a part of those generated in *P. tricornutum*. Additional metabolites could have been observed using different sample preparation or may have been extracted but remained undetected due to low ionization efficiency.

DCF metabolites previously observed in photosynthetic organisms such as 1-(2,6-dichlorophenyl)-5hydroxy-2-indolinone and acyl-glutamatyl-diclofenac (Fu et al., 2017; Liakh et al., 2023) were not detected in *P. tricornutum* in the present study. Sugars can be involved in the formation of metabolites in diatoms, for instance glucoronide conjugated metabolites were reported in *Navicula* sp. exposed to triclosan (Ding et al., 2018). However, the large size (m/z > 560) of the Phase II metabolites detected in the present study exclude the conjugation with sugars. Observed metabolites may instead be formed upon conjugation with amino acids or dipeptides. Synthesis and accumulation of Aa and proteins was reported in phytoplankton exposed to contaminants (Sharma et al., 2023; Zhang et al., 2021) and amino acid conjugates were observed in diatoms biotransformation pathways (Ding et al., 2018). In the present work, gene ontology analysis indicated that Aa and peptide biosynthetic pathways were regulated upon DCF exposure supporting a possible correlation between organic contaminants detoxification responses, amino acid and protein synthesis.

Treatment with CYP inhibitor fluconazole induced a change in the abundance of DCF and in two of its metabolites (M3 and M7). DCF intracellular concentrations were significantly higher in cells treated with fluconazole indicating that its biotransformation is affected. Furthermore, the abundance of metabolite M7 decreased under this treatment and amounts of M3 increased upon fluconazole treatment suggesting that cells may use alternative transformation pathways to compensate for the inhibition of one of them.

## 5. Conclusion

The present work investigated in parallel physiological effects, gene expression and biotransformation pathways associated with DCF exposure in the marine diatom *P. tricornutum*.

DCF resulted in mild physiological effects on *P. tricornutum* but gene expression analysis indicated that multiple molecular functions and biological processes were altered by DCF exposure. Genes involved in photosynthesis (light reaction and light harvesting), lipid, Aa and peptides biosynthesis were up regulated together with genes involved in iron homeostasis. Transcriptomic analysis suggested increased nutrients and energy requirements possibly associated with the contaminant stress and detoxification metabolism.

Removal of DCF from exposure medium was not observed, indicating that this marine diatom is not expected to be competitive with respect to freshwater green algae in DCF bio sequestration biotechnological processes. DCF intracellular concentrations were found to be relatively low with respect to DCF exposure concentrations suggesting that DCF may be internalized at low rates and/or that *P. tricornutum* is able to activate effective biotransformation pathways. Multiple phase II metabolites with high molecular weight were detected in *P. tricornutum* cells together with OH-DCF. The hydroxylation reaction associated to the formation of this metabolite is generally associated to CYP enzymatic activity. However, CYP genes were not significantly upregulated upon DCF exposure. On the other hand, two genes encoding for enzymes with cytochrome B5 domain were up regulated in a concentration dependent manner suggesting a possible involvement of CYP redox partners in DCF biotransformation. Treatment with CYP inhibitor increased the intracellular DCF concentration and altered the abundance of metabolites. However, obtained results could not provide evidence for a clear involvement of CYP enzymes in DCF biotransformation. Multiple DCF biotransformation pathways may be activated in parallel. Investigation of such alternative pathways, identification of involved enzymes and their encoding genes would help better understand the diversity of transformation strategies in phytoplankton.

## Acknowledgements

The authors thank the Platform Of Non-Target Environmental Metabolomics (PONTEM) of the Montpellier Alliance for Metabolomics and Metabolism Analysis (MAMMA) consortium facilities. The authors thank Christophe Leboulanger for scientific discussion and for proofing the manuscript.

## Author contributions

CrediT

**L. Alezra**: Investigation, Formal analysis **E. Le Floch**: Resources, Investigation, Methodology, Data curation **C. Felix**: Resources, Investigation, Methodology **D. Rosain:** Methodology **E. Gomez**: Resources, Methodology, Validation, Writing – review & editing

**F. Courant**: Supervision, Resources, Software, Formal analysis, Validation, Writing – review & editing **E. Fouilland**: Funding acquisition, Project administration, Supervision, Resources, Writing – review & editing **G. Cheloni:** Funding acquisition, Project administration, Supervision, Resources, Conceptualization, Investigation, Methodology, Data curation, Formal analysis, Validation, Visualization, Writing – original draft, Writing – review & editing

**Funding sources:** The work was developed during PHYCOCYP project. This project has received funding from the European Union’s Horizon 2020 research and innovation programme under the Marie Sklodowska-Curie grant agreement No 101030396

## Notes

### Competing Interest Statement

The authors have declared no competing interest.

