## Supplemental Material for "Diclofenac stress responses and biotransformation pathways in the marine diatom *Phaeodactylum tricornutum*"

Number of pages (including this page): 9
Number of Figures: 6
Number of Tables: 3

<u>Growth of *P.tricornutum* cultures</u>

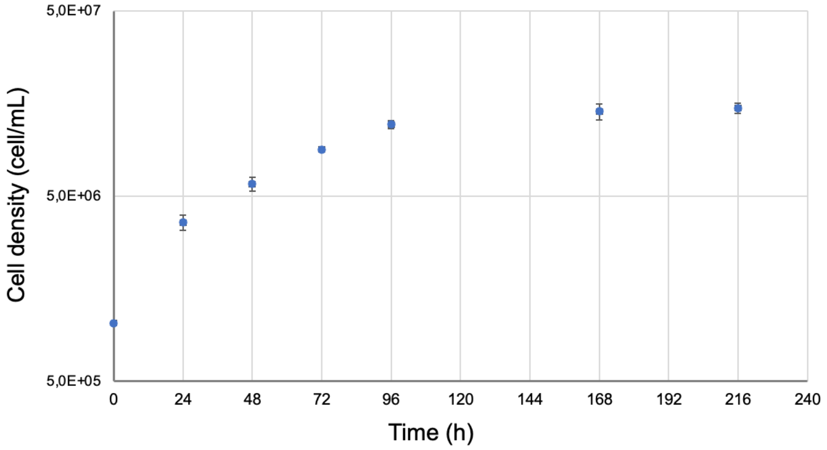


**Figure S1**: Growth curve of *P. tricornutum* CCAP1055/18 cultures grown in f/2 medium under controlled conditions (20°C, continuous agitation at120 rpm, warm white fluorescent lighting at 50 µE with a 16/8 light:dark cycle)

### Axenicity of *P.tricornutum* cultures

*P. tricornutum* CCAP1055/18 cultures axenicity was investigated in the beginning and at the end of the diclofenac exposure tests. Samples of cultures were collected and stained with the SYTO BC bacteria stain (Bacteria counting kit, Molecular Probes, Invitrogen) following the manufacturer protocol. Samples were analyzed using the flow cytometer BD Accuri C6 plus (BD biosciences). Gating strategy adopted to quantify diatom cells and bacterial cells is described in Figure S2.

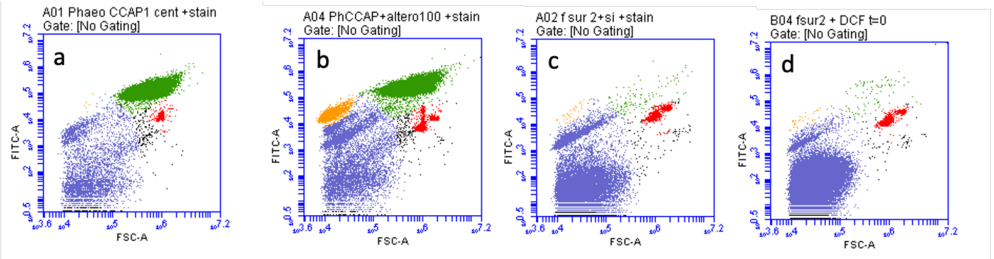


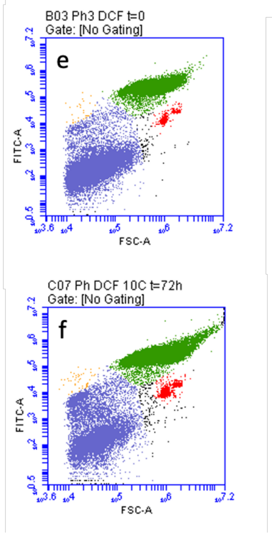


| Sample | Normalized count diatom | Normalized count bacteria | Bacteria/diatom ratio |
|---|---|---|---|
| Inoculum A | 1386 | 7 | 0,005 |
| Inoculum B | 1677 | 5 | 0,003 |
| Inoculum C | 2060 | 4 | 0,002 |
| C- A t=72h | 4622 | 71 | 0,015 |
| 1.5mg/L A DCF t=72h | 3474 | 6 | 0,002 |
| 10 mg/L A DCF t=72h | 3106 | 5 | 0,002 |
| C- B t=72h | 3564 | 23 | 0,007 |
| 1.5mg/L B DCF t=72h | 3366 | 15 | 0,005 |
| 10 mg/L B DCF t=72h | 2481 | 13 | 0,005 |
| C- C t=72h | 5842 | 45 | 0,008 |
| 1.5mg/L C DCF t=72h | 2178 | 5 | 0,002 |
| 10 mg/L C DCF t=72h | 3024 | 4 | 0,001 |
| f/2+Si+DCF | 3 | 1 | 0,333 |

**Figure S2**: Verification of algae axenicity before and during the exposure test. Gates for quantification of algal cells (green population), bacterial cells (yellow population) and inert particles (violet population) were defined analyzing axenic *P. tricornutum* cultures (a), *P. tricornutum* cultures inoculated with *Alteromonas* cells (b), f/2+Si medium (c) and f/2+Si medium with diclofenac at 10 mg $L^{-1}$ concentration (d). E xamples of cytograms obtained for samples collected at the beginning of the test (t=0) and at the end of the test (t=72h) are reported in € and (f) respectively. The table displays the counts of diatom and bacterial cells determined in each sample and normalized with respect to the beads (red population). The number of bacterial cells measured in the test cultures was comparable to the noise measured in the sterile exposure medium and did not change during the test. Even assuming the presence of bacterial cells inside *P. tricornutum* cultures, their number would be more than two orders of magnitude lower than diatom cells and can be considered irrelevant with respect to the obtained results.

#### HPLC sequence and procedure used to measure DCF in exposure medium

Injection volume was set at 20 µL with a 1 mL/min flow rate. The working UV wavelength was 220 nm. For DCF samples concentration was quantified based on calibration curved that were obtained for different concentration points. Two different calibration curves were used. The curve for lower DCF concentrations included 5 calibration points (0.05, 0.1, 0.5, 1 and 2 mg/L). The curve for higher DCF concentrations included 5 additional calibration points 5 calibration points (1, 7, 15, 30 and 60 mg/L). Calibration standards were prepared from a stock solution of DCF dissolved in EtOH. During sample measurement, Instrument stability was checked by regularly injecting a concentration corresponding to one point on the calibration curve during the analysis sequence. Peak integration and data analyses were performed using the Chromoleon software. Measurements were performed in 10 × concentrated SPE extracts. The concentrations in f/2 medium samples were calculated taking the extraction recoveries into account.

A

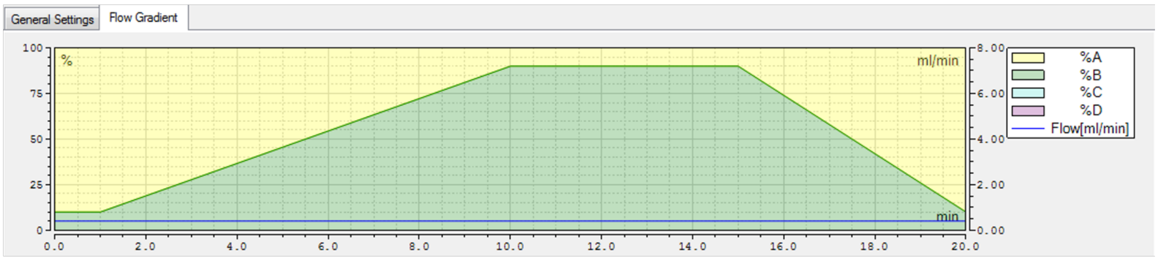


B

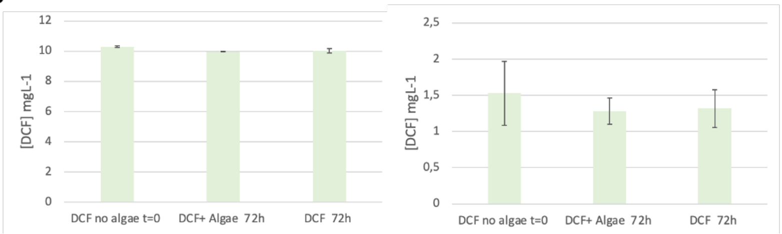


**Figure S3**: (A) HPLC-UV gradient used for DCF determination in exposure medium, phase A corresponds to water, phase B corresponds to acetonitrile. (B) DCF concentrations measured in exposure medium in the beginning and at the end of the test in presence and absence of algae.

### Flow cytometry measurements of intracellular oxidative stress

Detection of reactive oxygen species (ROS) in living cells was performed via flow cytometry using the CellROX™ green flow cytometry assay kit (Molecular Probes, Life Technologies). Samples were collected after 24, 48 and 72 hours from control and exposed cultures (1.5 and 10 mg $L^{-1}$ DCF) and stained following the manufacturer protocol. Samples were analyzed using the flow cytometer BD Accuri C6 plus (BD biosciences). Gating strategy was designed based on negative and positive controls. Two gates were identified. The Cell Rox (CR) positive gate was designed to detect populations of cells undergoing oxidative stress that was induced using the tert-butyl hydroperoxide solution (TBHP, an ROS inducer) provided in the kit. The Cell Rox (CR) negative gate was designed to detect populations of cells that were not affected by oxidative stress, these cells were treated with N-acetylcysteine (NAC) to increase the antioxidant capability of the cell before performing the tert-butyl hydroperoxide treatment. Detailed information about solutions concentrations and incubation time can be found in the manufacturer protocol.

A

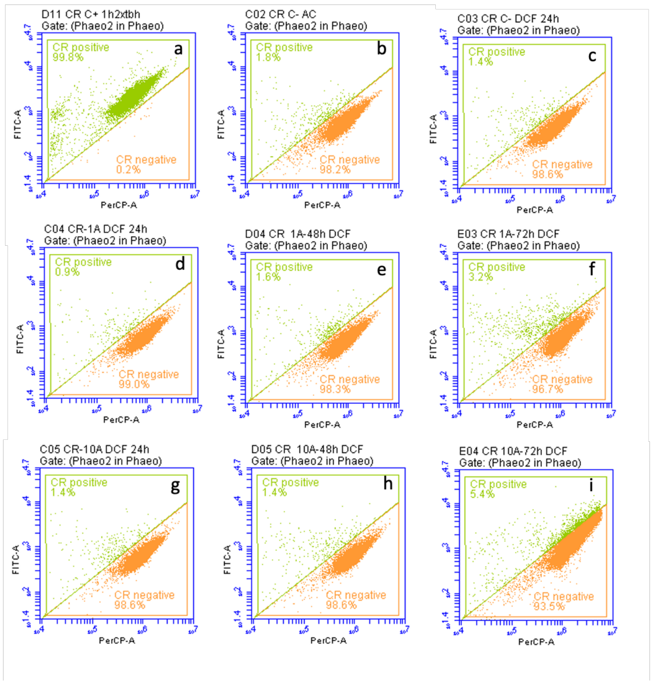


B

| Samples | CR negative | CR positive |
|---|---|---|
| C03 CR C- DCF 24h | 98.57% | 1.40% |
| F02 CR C- t=24 | 96.88% | 3.07% |
| F02 CR C- 24h | 98.28% | 1.65% |
| C04 CR-1A DCF 24h | 99.02% | 0.94% |
| C06 CR-1B DCF 24h | 98.73% | 1.24% |
| C08 CR-1C DCF 24h | 98.31% | 1.68% |
| C05 CR-10A DCF 24h | 98.62% | 1.35% |
| C07 CR-10B DCF 24h | 98.57% | 1.43% |
| C09 CR-10C DCF 24h | 98.71% | 1.26% |
| D02 CR C-48 | 98.54% | 1.43% |
| G02 CR C- 48h | 97.07% | 2.81% |
| G02 CR C- 48 | 98.99% | 0.94% |
| D04 CR 1A-48h DCF | 98.34% | 1.62% |
| D06 CR 1B-48h DCF | 98.88% | 1.08% |
| D08 CR 1C-48h DCF | 98.92% | 1.03% |
| D05 CR 10A-48h DCF | 98.59% | 1.40% |
| D07 CR 10B-48h DCF | 98.50% | 1.47% |
| D09 CR 10C-48h DCF | 98.80% | 1.14% |
| E02 CR C-72h DCF | 98.44% | 1.52% |
| H02 CR C- 72h | 96.58% | 3.16% |
| H02 CR C -72h | 94.19% | 2.33% |
| E03 CR 1A-72h DCF | 96.65% | 3.22% |
| E05 CR 1B-72h DCF | 97.11% | 2.83% |
| E07 CR 1C-72h DCF | 97.32% | 2.24% |
| E04 CR 10A-72h DCF | 93.50% | 5.35% |
| E06 CR 10B-72h DCF | 96.96% | 2.09% |
| E08 CR 10C-72h DCF | 97.35% | 1.64% |

**Figure S4**: Effects of Diclofenac on intracellular oxidative stress. **A**) Examples of cytograms obtained for positive control (TBHP treatment) (a), negative control (NAC+TBHP treatment) (b) and not exposed control culture of the Diclofenac treatment experiment (c). ROS in cells treated with DCF 1.5 $mgL^{-1}$ (d, e, f) and 10 mg $L^{-1}$ (g, h, l) were measured after 24, 48 and 72 hours incubation. **B**) Table with measured percentages of not affected (CR negative) and affected (CR positive) cells obtained for the three treatments (control, DCF1.5 $mgL^{-1}$ and 10 mg $L^{-1}$) after 24, 48 and 72 hours incubation.

**Table S1**: List of genes differentially expressed in 10 mg $L^{-1}$ treatment with respect to 1.5 mg $L^{-1}$ treatment that have a DNA binding function.

| Term ID | Term name | Adjusted p value | intersection size | Gene ID | Transcript ID | Gene Symbol | **10 mg/1.5 mg** |
|---|---|---|---|---|---|---|---|
| GO:0003700 | DNA-binding transcription factor activity | **0,019581495** | 30 | 7195220 | XM_002183504 | DEL | **5,027548** |
| GO:0003700 | DNA-binding transcription factor activity | **0,019581495** | 30 | 7195452 | XM_002183737 | . | **-2,705821** |
| GO:0003700 | DNA-binding transcription factor activity | **0,019581495** | 30 | 7195641 | XM_002183760 | . | **-3,462452** |
| GO:0003700 | DNA-binding transcription factor activity | **0,019581495** | 30 | 7195642 | XM_002183761 | . | **-3,090048** |
| GO:0003700 | DNA-binding transcription factor activity | **0,019581495** | 30 | 7195662 | XM_002183781 | . | **-3,569197** |
| GO:0003700 | DNA-binding transcription factor activity | **0,019581495** | 30 | 7196246 | XM_002177362 | . | **-5,719785** |
| GO:0003700 | DNA-binding transcription factor activity | **0,019581495** | 30 | 7196272 | XM_002176556 | . | **-4,526316** |
| GO:0003700 | DNA-binding transcription factor activity | **0,019581495** | 30 | 7197371 | XM_002177514 | . | **-2,217568** |
| GO:0003700 | DNA-binding transcription factor activity | **0,019581495** | 30 | 7197429 | XM_002177573 | . | **-2,163617** |
| GO:0003700 | DNA-binding transcription factor activity | **0,019581495** | 30 | 7197512 | XM_002178042 | . | **-2,403998** |
| GO:0003700 | DNA-binding transcription factor activity | **0,019581495** | 30 | 7198088 | XM_002178576 | . | **-2,046734** |
| GO:0003700 | DNA-binding transcription factor activity | **0,019581495** | 30 | 7198226 | XM_002184347 | . | **-2,339921** |
| GO:0003700 | DNA-binding transcription factor activity | **0,019581495** | 30 | 7198249 | XM_002184368 | . | **-4,903104** |
| GO:0003700 | DNA-binding transcription factor activity | **0,019581495** | 30 | 7198250 | XM_002184371 | HSF2 | **-4,430202** |
| GO:0003700 | DNA-binding transcription factor activity | **0,019581495** | 30 | 7198282 | XM_002184388 | . | **-2,771365** |
| GO:0003700 | DNA-binding transcription factor activity | **0,019581495** | 30 | 7198798 | XM_002184883 | . | **-2,566059** |
| GO:0003700 | DNA-binding transcription factor activity | **0,019581495** | 30 | 7199608 | XM_002178781 | . | **-2,239250** |
| GO:0003700 | DNA-binding transcription factor activity | **0,019581495** | 30 | 7200092 | XM_002179404 | . | **-2,041418** |
| GO:0003700 | DNA-binding transcription factor activity | **0,019581495** | 30 | 7200184 | XM_002179126 | . | **-5,198998** |
| GO:0003700 | DNA-binding transcription factor activity | **0,019581495** | 30 | 7200308 | XM_002179351 | . | **-5,822985** |
| GO:0003700 | DNA-binding transcription factor activity | **0,019581495** | 30 | 7200327 | XM_002179145 | . | **-5,831303** |
| GO:0003700 | DNA-binding transcription factor activity | **0,019581495** | 30 | 7200452 | XM_002179894 | . | **-2,098044** |
| GO:0003700 | DNA-binding transcription factor activity | **0,019581495** | 30 | 7200486 | XM_002179735 | . | **-7,733135** |
| GO:0003700 | DNA-binding transcription factor activity | **0,019581495** | 30 | 7200625 | XM_002179829 | . | **-7,329756** |
| GO:0003700 | DNA-binding transcription factor activity | **0,019581495** | 30 | 7201958 | XM_002181395 | . | **4,692164** |
| GO:0003700 | DNA-binding transcription factor activity | **0,019581495** | 30 | 7202319 | XM_002181462 | . | **-4,348011** |
| GO:0003700 | DNA-binding transcription factor activity | **0,019581495** | 30 | 7202512 | XM_002181681 | . | **-5,261197** |
| GO:0003700 | DNA-binding transcription factor activity | **0,019581495** | 30 | 7203194 | XM_002182372 | . | **-2,634361** |
| GO:0003700 | DNA-binding transcription factor activity | **0,019581495** | 30 | 7203588 | XM_002182779 | . | **-3,276483** |
| GO:0003700 | DNA-binding transcription factor activity | **0,019581495** | 30 | 7204220 | XM_002186084 | . | **-2,060471** |

**Table S2**: List of genes differentially expressed in 1.5 mg $L^{-1}$ treatment and 10 mg $L^{-1}$ treatment with respect to control and that bind a tetrapyrrole. Genes are grouped between Chloropyll binding (upper part of the table) and Heme binding (lower part of the table).

| Gene_ID | Name_locusTag | name | description | Binding | FC DCF 1.5 mgL-1/C- | FC DCF 10 mgL-1/C- |
|---|---|---|---|---|---|---|
| 7195106 | . PHATRDRAFT_48798 | - | Fucoxanthin chlorophyll a/c protein, deviant | Chl | 1.752975 | 2.212798 |
| 7195163 | Lhcf15 PHATRDRAFT_48882 | LHCF15 | Protein fucoxanthin chlorophyll a/c protein | Chl | 7.899762 | 10.675412 |
| 7195300 | Lhcr2 PHATRDRAFT_22956 | LHCR2 | Protein fucoxanthin chlorophyll a/c protein | Chl | 1.335797 | 2.112161 |
| 7195835 | Lhcr11 PHATRDRAFT_23257 | LHCR11 | Protein fucoxanthin chl a/c protein | Chl | 1.856068 | 2.88852 |
| 7196166 | . PHATRDRAFT_17531 | - | Fucoxanthin chlorophyll a/c protein | Chl | 2.318131 | 2.759127 |
| 7196834 | Lhcr12 PHATRDRAFT_54027 | LHCR12 | Protein fucoxanthin chlorophyll a/c protein | Chl | 2.027724 | 2.254238 |
| 7196903 | Lhcr8 PHATRDRAFT_32294 | LHCR8 | Protein fucoxanthin chlorophyll a/c protein | Chl | 4.366406 | 1.299127 |
| 7197205 | Lhcr7 PHATRDRAFT_43522 | LHCR7 | Protein fucoxanthin chlorophyll a/c protein | Chl | 4.153307 | 1.497715 |
| 7198473 | Lhcf7 PHATRDRAFT_30643 | LHCF6 | Protein fucoxanthin chlorophyll a/c protein | Chl | 2.092204 | 3.316313 |
| 7198609 | Lhcf17 PHATRDRAFT_56310 | LHCF17 | Protein fucoxanthin chlorophyll a/c protein | Chl | 1.339034 | 2.635791 |
| 7198763 | Lhcr10 PHATRDRAFT_50086 | LHCR10 | Protein fucoxanthin chlorophyll a/c protein | Chl | -2.441368 | -2.048021 |
| 7199273 | . PHATRDRAFT_24119 | - | Fucoxanthin chlorophyll a/c protein, deviant | Chl | 1.660467 | 5.453404 |
| 7199648 | Lhcf16 PHATRDRAFT_34536 | LHCF16 | Protein fucoxanthin chlorophyll a/c protein | Chl | 2.503917 | 5.242158 |
| 7199712 | Lhcx3 PHATRDRAFT_44733 | LHCX3 | Protein fucoxanthin chlorophyll a/c protein | Chl | 2.118705 | -1.364067 |
| 7200476 | Lhcx1 PHATRDRAFT_27278 | LHCX1 | Protein fucoxanthin chlorophyll a/c protein | Chl | 4.262509 | 4.864009 |
| 7202270 | . PHATRDRAFT_47485 | - | Fucoxanthin chlorophyll a/c protein, deviant | Chl | 1.440629 | 2.670683 |
| 7202932 | Lhcr6 PHATRDRAFT_56319 | LHCR6 | Protein fucoxanthin chlorophyll a/c protein | Chl | 3.434936 | 1.322108 |
| 7202961 | Lhcr13 PHATRDRAFT_38121 | LHCR13 | Protein fucoxanthin chlorophyll a/c protein | Chl | 1.467414 | 2.418821 |
| 7203055 | Lhcr14 PHATRDRAFT_47813 | LHCR14 | Protein fucoxanthin chlorophyll a/c protein | Chl | 2.014963 | 4.033398 |
| 7203170 | Lhcf10 PHATRDRAFT_22006 | LHCF10 | Protein fucoxanthin chlorophyll a/c protein | Chl | 1.639624 | 2.577897 |
| 7203256 | Lhcf6 PHATRDRAFT_29266 | LHCF6 | Protein fucoxanthin chlorophyll a/c protein | Chl | 2.196949 | 3.379733 |
| 7203558 | . PHATRDRAFT_6062 | - | LHC 6062 | Chl | 2.352313 | 4.348967 |
| 7203582 | Lhcf8 PHATRDRAFT_22395 | LHCF8 | Protein fucoxanthin chlorophyll a/c protein | Chl | 2.868779 | 4.206019 |
| 7203779 | Lhcx4 PHATRDRAFT_38720 | LHCX4 | Protein fucoxanthin chlorophyll a/c protein | Chl | -1.176121 | -3.264915 |
| 7204099 | Lhcf14 PHATR_25893 | LHCF14 | Fucoxanthin chlorophyll a/c protein, lhcf type | Chl | 2.113743 | 3.448886 |
| 7195268 | . PHATRDRAFT_30040 | EG02251 | Cyt_B5-like_heme binding | Heme | -1.623065 | -2.310334 |
| 7195287 | . PHATRDRAFT_54983 | - | Nitrate reductase | Heme | 1.031158 | 3.002762 |
| 7195613 | . PHATRDRAFT_39797 | - | Heme binding | Heme | -2.187038 | -2.813575 |
| 7197527 | . PHATRDRAFT_43466 | - | CYP5165A2 | Heme | 3.154697 | 1.482711 |
| 7197528 | . PHATRDRAFT_43467 | - | CYP5165A1 | Heme | -1.508719 | -2.114846 |
| 7198555 | . PHATRDRAFT_30770 | - | Cyt_B5_heme-binding ER localized | Heme | -1.075005 | 2.502166 |
| 7199210 | . PHATRDRAFT_8141 | - | heme binding | Heme | 1.674423 | 2.421983 |
| 7199255 | . PHATRDRAFT_50443 | EG02619 | Fatty acid desaturase with Cyt_B5-like_ domain | Heme | 2.709718 | 7.550731 |
| 7199710 | . PHATRDRAFT_51720 | - | Fumarate reductase flavoprotein Cyt_B5-like domain | Heme | 1.339314 | -2.081438 |
| 7200392 | . PHATRDRAFT_45621 | - | NOD nitric oxide dioxygenase | Heme | -1.404601 | -5.965809 |
| 7200407 | . PHATRDRAFT_35395 | - | Heme binding with Cyt_B5-like_domain | Heme | 5.333752 | 9.029313 |
| 7200713 | . PHATRDRAFT_19954 | - | Heme binding | Heme | -1.75459 | -3.777496 |
| 7201461 | . PHATRDRAFT_27757 | - | NiR-Fd. ferredoxin-nitrite reductase | Heme | 1.818732 | 3.854727 |
| 7201607 | . PHATRDRAFT_13174 | EG02288 | Ascorbate peroxidase | Heme | 1.521744 | 4.674537 |
| 7201664 | . PHATRDRAFT_8155 | EG02286 | Heme binding | Heme | -4.07866 | -2.485417 |
| 7202394 | . PHATRDRAFT_47332 | - | Cyt-C6 | Heme | -1.496248 | -2.547627 |
| 7202539 | . PHATRDRAFT_47395 | - | peroxidase activity | Heme | 2.233298 | 1.655188 |
| 7203399 | . PHATRDRAFT_48278 | - | Oxidoreductase activity | Heme | 4.87577 | 1.910832 |
| 7203402 | . PHATRDRAFT_48286 | - | Peroxidase activity | Heme | 4.807917 | 1.903368 |
| 7203792 | . PHATRDRAFT_29488 | D6 | Delta 6 fatty acid desaturase (Cyt_B5-like_heme) | Heme | 1.360202 | 2.441332 |
| 7203920 | . PHATR_33568 | - | Heme binding | Heme | 2.302948 | -1.119226 |
| 7204234 | ]. PHATR_44056 | PETJ | Cytochrome c6, cytochrome c553 | Heme | 2.046366 | 2.004875 |
| 7204578 | . PHATR_20948 | - | Oxidoreductase activity | Heme | 2.045373 | 1.608874 |
| 7204592 | . PHATR_13244 | - | Cytochrome *c* peroxidase | Heme | 2.678183 | 2.291865 |
| 7204684 | . PHATR_46830 | PTD5A | Delta 5 fatty acid desaturase (Cyt_B5-like_ domain) | Heme | 1.507082 | 2.442625 |

**Table S3**: List of top 15 genes significantly up regulated in the two DCF treatments with respect to control

| Gene_ID | Transcript_ID | Gene_Symbo | Description | gene_biotype | Protein_ID | locus_tag | **1.5 mgL/C-.fc** |
|---|---|---|---|---|---|---|---|
| 7195419 | XM_002183706 | **ISIP2B** | iron starvation induced protein | protein_coding | XP_002183742.1 | PHATRDRAFT_54987 | **79,069279** |
| 7200478 | XM_002179726 | **ISIP2A** | iron starvation induced protein | protein_coding | XP_002179762.1 | PHATRDRAFT_54465 | **38,519051** |
| 7195295 | **XM_002183568** | . | **cell surface protein** | protein_coding | XP_002183604.1 | PHATRDRAFT_54986 | **36,374577** |
| 7195420 | XM_002183707 | **CREG1** | cellular repressor of e1a-stimulated gene-like protein | protein_coding | XP_002183743.1 | PHATRDRAFT_51183 | **35,807480** |
| 7203427 | XM_002182582 | . | predicted protein | protein_coding | XP_002182618.1 | PHATRDRAFT_22166 | **29,539665** |
| 7198713 | XM_002184863 | . | flavodoxin | protein_coding | XP_002184899.1 | PHATRDRAFT_23658 | **23,732582** |
| 7199186 | XM_002185288 | . | predicted protein | protein_coding | XP_002185324.1 | PHATRDRAFT_50361 | **21,841268** |
| 7195234 | XM_002183518 | . | predicted protein | protein_coding | XP_002183554.1 | PHATRDRAFT_39557 | **20,495379** |
| 7199595 | XM_002178986 | . | predicted protein | protein_coding | XP_002179022.1 | PHATRDRAFT_34433 | **20,360843** |
| 7195296 | **XM_002183569** | . | **cell surface protein** | protein_coding | XP_002183605.1 | PHATRDRAFT_52498 | **19,569704** |
| 7202689 | XM_002182039 | **ISIP3** | iron starvation induced protein | protein_coding | XP_002182075.1 | PHATRDRAFT_47674 | **17,929683** |
| 7196953 | XM_002177470 | . | predicted protein | protein_coding | XP_002177506.1 | PHATRDRAFT_43232 | **16,959829** |
| 7194827 | XM_002183052 | . | predicted protein | protein_coding | XP_002183088.1 | PHATRDRAFT_48621 | **16,757517** |
| 7201479 | XM_002180478 | . | predicted protein | protein_coding | XP_002180514.1 | PHATRDRAFT_46275 | **14,234892** |
| 7195800 | XM_002184057 | . | predicted protein | protein_coding | XP_002184093.1 | PHATRDRAFT_40136 | **13,327611** |

| Gene_ID | Transcript_ID | Gene_Symbo | Description | gene_biotype | Protein_ID | locus_tag | **10 mgL/C-.fc** |
|---|---|---|---|---|---|---|---|
| 7195419 | XM_002183706 | **ISIP2B** | iron starvation induced protein | protein_coding | XP_002183742.1 | PHATRDRAFT_54987 | **550,881607** |
| 7195295 | **XM_002183568** | . | **cell surface protein** | protein_coding | XP_002183604.1 | PHATRDRAFT_54986 | **245,072425** |
| 7195420 | XM_002183707 | **CREG1** | cellular repressor of e1a-stimulated gene-like protein | protein_coding | XP_002183743.1 | PHATRDRAFT_51183 | **212,500794** |
| 7200478 | XM_002179726 | **ISIP2A** | iron starvation induced protein | protein_coding | XP_002179762.1 | PHATRDRAFT_54465 | **103,602583** |
| 7195296 | **XM_002183569** | . | **cell surface protein** | protein_coding | XP_002183605.1 | PHATRDRAFT_52498 | **92,564529** |
| 7198713 | XM_002184863 | . | flavodoxin | protein_coding | XP_002184899.1 | PHATRDRAFT_23658 | **76,805300** |
| 7203427 | XM_002182582 | . | predicted protein | protein_coding | XP_002182618.1 | PHATRDRAFT_22166 | **27,383038** |
| 7199266 | XM_002185395 | FbaC5 | fructose-bisphosphate aldolase | protein_coding | XP_002185431.1 | PHATRDRAFT_51289 | **26,081052** |
| 7196604 | XM_002176951 | Lhcx2 | protein fucoxanthin chlorophyll a/c protein | protein_coding | XP_002176987.1 | PHATRDRAFT_54065 | **17,896176** |
| 7196331 | XM_002177121 | . | predicted protein | protein_coding | XP_002177157.1 | PHATRDRAFT_53967 | **17,890521** |
| 7202689 | XM_002182039 | **ISIP3** | iron starvation induced protein | protein_coding | XP_002182075.1 | PHATRDRAFT_47674 | **14,649909** |
| 7196953 | XM_002177470 | . | predicted protein | protein_coding | XP_002177506.1 | PHATRDRAFT_43232 | **14,439289** |
| 7195234 | XM_002183518 | . | predicted protein | protein_coding | XP_002183554.1 | PHATRDRAFT_39557 | **12,070122** |
| 7197243 | XM_002178067 | . | triosephosphate isomerase | protein_coding | XP_002178103.1 | PHATRDRAFT_50738 | **11,062869** |
| 7195563 | **XM_002183835** | . | **cell surface protein** | protein_coding | XP_002183871.1 | PHATRDRAFT_49272 | **10,885047** |

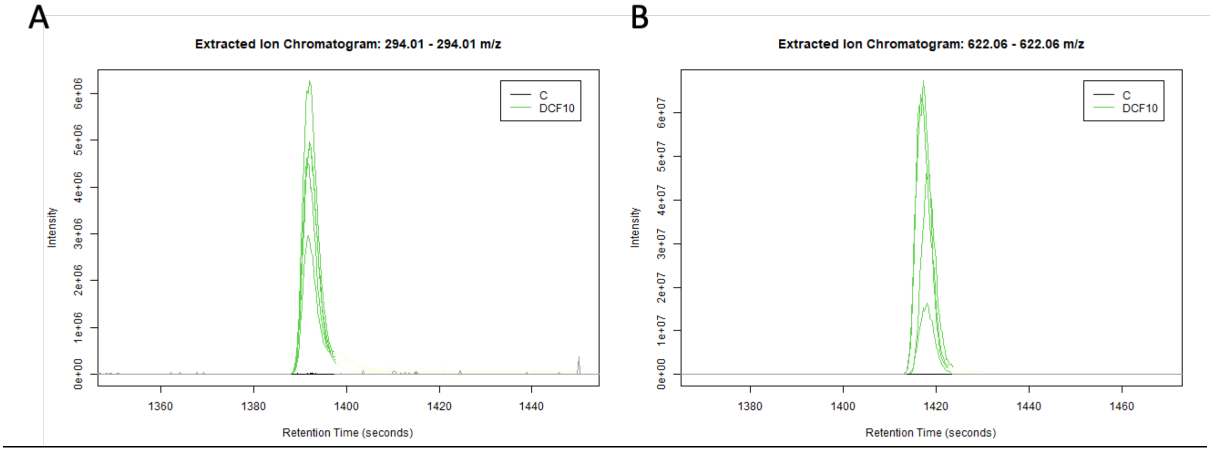


**Figure S5**: Peaks obtained with HPLC-Orbitrap MS analysis of cultures exposed to 10 mg $L^{-1}$ DCF. (A) peak of DCF and (B) peak of metabolite M6

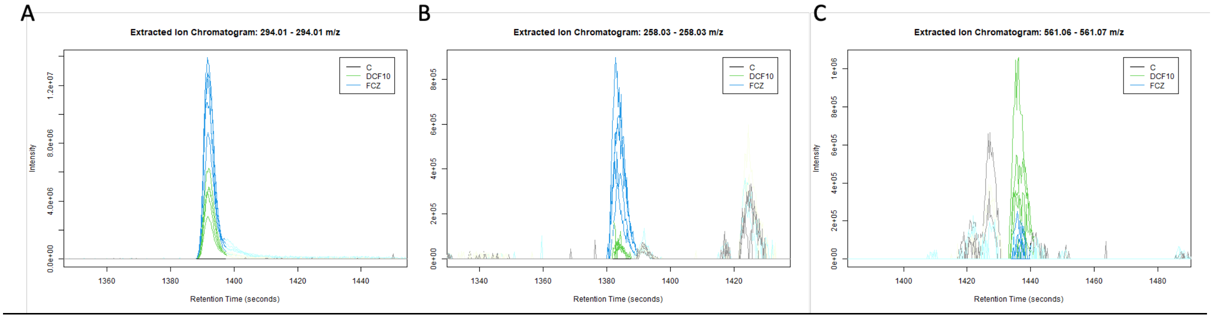


**Figure S6**: Peaks obtained with HPLC-Orbitrap MS analysis of cultures exposed to 10 mg $L^{-1}$ DCF (green line) and to 10 mg $L^{-1}$ DCF+Fluconazole (blue line) (A) peak of DCF, (B) peak of metabolite M3, (C) peak of metabolite M7.
